# Robust single-trial estimates of electrocortical generalized aversive conditioning: Validation of a Bayesian multilevel learning model

**DOI:** 10.1101/2025.02.16.638555

**Authors:** Andrew H. Farkas, Judith Cediel Escobar, Faith E. Gilbert, Mingzhou Ding, Andreas Keil

## Abstract

Aversive conditioning changes visuocortical responses to conditioned cues, and the generalization of these changes to perceptually similar cues may provide mechanistic insights into anxiety and fear disorders. Yet, neuroimaging conditioning paradigms are challenged by poor single-trial signal-to-noise ratios (SNR), missing trials, and inter-individual differences in learning. Here, we address these issues with the validation of a steady-state visual evoked potential (ssVEP) generalization paradigm in conjunction with a Rescorla-Wagner inspired Bayesian multilevel learning model. A preliminary group of observers (N=24) viewed circular gratings varying in grating orientation, with only one orientation paired with an aversive outcome (noxious electric pulse). Gratings were flickered at 15 Hz to evoke ssVEPs recorded with 31 channels of EEG in an MRI scanner. The multilevel structure of the Bayesian model learning model informs and constrains estimates per participant providing an interpretable generative model. It led to superior cross-validation accuracy and insights into individual participant dynamics than simpler models. It also isolates the generalized effects of conditioning, providing improved statistical certainty. Lastly, the present report demonstrates that missing trials are interpolated and weighted appropriately using the full model’s structure. This is a critical aspect for single-trial analyses of simultaneously recorded physiological measures, because each added measure will typically increase the number of trials missing a complete set of observations. The present technical report validates a limited version of a learning model to illustrate the utility of this analytical framework. It shows how models may be iteratively built and compared in a modern Bayesian workflow. Future models may use different conceptualizations of learning, allow integration of clinically relevant factors, and enable the fusion of different simultaneous physiological recordings.

## Introduction

Experience changes the tuning of visuocortical neurons, heightening the sensitivity to simple features that selectively predict motivationally relevant outcomes (Li & Keil, 2023; Shuler & Bear, 2006). This process is readily studied by means of Pavlovian generalization conditioning (Lissek et al., 2014). In this paradigm, stimuli varying in similarity are presented, but only one stimulus (the CS+) is consistently paired with the unconditioned stimulus (US, e.g., loud noise, electric shock), whereas other stimuli (generalization stimuli, GSs) are never paired. A test phase tends to follow an initial conditioning phase (Lonsdorf et al., 2017). Typical findings include that response variables such as skin conductance or pupil diameter are heightened when viewing the CS+, and that this enhancement is partly transferred to the GSs, as a function of their similarity with the CS+ (Ahrens et al., 2016). Neuroimaging studies of aversive generalization conditioning have found similar activation patterns in brain structures such as the amygdala or anterior insula (Dunsmoor & Murphy, 2015; Lissek et al., 2014). Studies interested in inter-individual differences have further observed that relatively heightened responses to GSs, referred to as overgeneralization, are often associated with elevated anxiety and negative affect (e.g., Lissek et al., 2008). Together, these observations highlight the foundational role of generalization learning for adaptive behavior.

What is currently not understood is how generalization learning emerges over time and how it shapes different physiological systems as observers learn to predict the aversive outcome. The present report illustrates the usage of Bayesian multilevel models in combination with steady-state visually evoked potentials (ssVEPs; see Norcia et al., 2015; Wieser et al., 2016) to index how aversive learning emerges in the visual brain, on a trial-by-trial basis. The ssVEP is evoked when a visual stimulus is flickered at a specific rate (here: 15 Hz), eliciting a strong electrocortical signal in sensors over visual cortex that oscillates at the same rate. Readily recorded from sensors placed over the scalp, the amplitude of electrical activity at the target frequency can be quantified in the frequency domain, being most pronounced at occipital recording sites. The ssVEP originates primarily in calcarine and peri-calcarine (lower-tier) visual areas. Over the past decades, we and others have established that the amplitude of the ssVEP varies with selective attention to motivationally relevant stimuli and that this modulation originates in visual and fronto-parietal cortical regions (Wieser & Keil, 2011, 2020). The ssVEP signal represents many periodic repetitions at a known specific frequency, and thus can be collapsed into one bin of a frequency-domain representation such as a Fourier spectrum, yielding a high signal-to-noise ratio (Muller & Hillyard, 2000; Petro et al., 2015; Regan, 1972). This property makes the ssVEP uniquely suitable for estimating robust single-trial amplitude values (Figueira et al., 2022). Such an ability is in contrast with most measures used in human neuroscience, which are dependent upon trial averaging to eliminate noise (Fabiani et al., 2007). A high signal-to-noise ratio is crucial for studies of learning in which dynamic trial-by-trial changes are expected.

However, regardless of the specific physiological measure used, missing trials and inter-individual differences complicate inference of dynamic processes during aversive conditioning. It is common for spontaneous artifacts to cause some trials to be unusable leading to “ragged” data sets with missing observations. This situation further deteriorates as more physiological measures are recorded simultaneously, leading to even less trials with a complete set of observations. Possibly more concerning is the difficulty to model inter-individual differences. It is expected that participants will learn aversive pairings at different rates, and aspects like generalization may depend on observer traits such as anxiety. Although this can be the explicit purpose and focus of a conditioning study, individual differences become another source of noise if not modeled correctly (Hedge et al., 2018). With traditional statistics, it is usually not feasible to model a moderate amount of complexity (like the learning rate per participant) without concerns related to statistical power and overfitting.

Bayesian multilevel models address problems in which granular estimates (e.g., individualized learning rates) are both informed and constrained by a hierarchal structure of priors (e.g., the average learning rate). The hierarchal structure also applies a by-default form of regularization, increasing cross-validation accuracy and correcting for multiple comparisons (Gelman et al., 2012). This allows for more features of the data to be modeled accurately and sensibly without overfitting concerns. A specific benefit for trial-dependent physiology studies is that missing trials can be specified as parameters to be estimated. This results in missing trials being interpolated not as a single point, but as a distribution informed by the full model’s structure and thus reflecting the appropriate amount of statistical uncertainty.

The aim of this technical report is to establish how a Bayesian multilevel learning model can be applied to single trial data from aversive conditioning paradigms. Three models are fit on a preliminary set of EEG-recorded ssVEP data (N = 24) in a differential conditioning paradigm recorded during fMRI recording. Each subsequent model builds in complexity, starting as a description of overall trends to the final model (Model 3) that estimates individual participant learning rates that use a version of the Rescorla-Wagner learning rule. In this algorithm, memory updating (learning) is quantified as a person’s prediction error on a given experimental trial, and the learning rate parameter (α) describes the speed of association acquisition. This model posits that as surprise increases, changes in associative strength are larger (Rescorla and Wagner, 1972; see eq. 1).

All three models compared in the present report interpolate missing trials and model inter-individual differences in the adaptation (reduction) of the ssVEP amplitude over the course of the study. After outlining EEG preprocessing and model equations, the models are compared in terms of cross-validation. Posterior distributions are then used to visualize and interpret the effects of interest which include skewed and bounded learning rates which would be difficult to fit and interpret with Frequentist multilevel models. Lastly, the Bayesian interpolation of missing trials is visualized. This latter feature is especially relevant for multimodal or multi-variable imaging studies, in which multiple measures (e.g. EEG, fMRI-BOLD, Heart Rate, Pupil diameter etc.) are recorded concurrently.

## 2 | Methods

### 2.1 | Participants

Participants were recruited from the University of Florida Psychology research pool and compensated with course credit. As this was a simultaneous EEG-fMRI paradigm, data collection was more complicated with more points of failure compared to EEG-only studies. 31 participants were run through the paradigm with EEG-recording. The first two participants were excluded because of data quality issues two due to experimenter error with new equipment. Another participant was excluded because of vision acuity issues limiting their ability to see the cues or read rating instructions. Two participants were excluded for excessive movement-related artifacts. The last exclusion was a participant that fell asleep during the paradigm. The final sample comprised of 24 participants, with 18 being Female and 8 identifying as Hispanic. The average age of the sample was 19.9 (*SD* = 1.78). The self-identified race of sample was 18 White, 3 Asian, 2 Multiracial, with one participant not reporting race. All participants gave written informed consent using forms approved by the University of Florida Institutional Review Board.

### 2.2 | Procedure

After arriving at the data collection building, participants were guided to a quiet preparatory room. After consenting, participants were then screened for the MRI scanning procedure to ensure no safety or medical concerns were present. After all the equipment was fit, but prior to entering the scanner, participants completed a series of questionnaires and demographic information.

Researchers then added Biopac spectra360 GEL104 to two EL509 MR-safe dry adhesive electrodes and applied the gelled electrodes to the participant’s left outer ankle. Electrodes were positioned two finger widths above the left ankle bone and slightly anterior to the fibula. The electrical stimulus was administered using the Biopac STMISOC system connected to the STM100C stimulator module, with LEAD108 connecting to the EL509 electrodes. A shock workup procedure was then implemented, where the electrical stimulus intensity was increased until participants stated that it was “unpleasant but not painful, and still tolerable”. This procedure was repeated twice. After obtaining the stimulus threshold, participants were then asked to rate a series of 5 shocks in row on a scale of 1 to 9, with 9 being extremely unpleasant, and 1 being not unpleasant at all. After the final shock, the participant was asked if the stimulus was still unpleasant but tolerable. The average shock level was 7.26 mA (SD = 5.0, min = 1 mA, max = 50 mA).

After the shock tuning, participants were fit with a 32-channel MR-compatible EEG system (BrainAmp MR; Brain Products). Gel was applied using blunt tip syringes and cotton swabs until impedance values were below 10kOhms or as low as possible in accordance with the Brain Product recommendations. The electrode for measuring ECG was placed on the participant’s back slightly to the left of the spine, while tucking the chin to allow enough slack for the cable while laying down in the scanner. A Siemens Head/Neck 64 coil was placed over the participant’s head, and the cable connecting the EEG to the amplifier was fed through the back of the head coil. A mirror attached to the head coil allowed participants to view the presentation monitor, which was located 70 inches away from the participant.

Participants were then led to the 3T chamber, and given earplugs to reduce any discomfort from the noise of the scanner. In their right hand, participants were given a clicker with a left and right button for making ratings throughout the study. After each block, participants were shown a Gabor pattern and were given six seconds to make their rating. They could click left and right to make their rating from the middle of a 11-point likert scale for ratings of hedonic valence and arousal. Additionally, participants were also asked to rate their expectancy of a shock for each pattern from 0 to 100% in 10% increments.

A two-way communication microphone was utilized to maintain communication between the participant and researcher for instructions and in case of emergency. Following the task, participants were removed from the chamber and asked to rate the unpleasantness of the shock the last time they felt it. Participants were also asked if they thought there was any relationship between the patterns and the electric stimulus. Only participant 10 was unable to state the correct conditioning relationship. After debriefing, participants were thanked and compensated for their time.

### 2.3 | Conditioning paradigm

This task involved a fear conditioning paradigm with a sequential design involving 4 blocks of Habituation, Acquisition #1, Acquisition #2, and Extinction. In this design, flickering Gabor patches with different grating orientations were presented for approximately 2 seconds with a random ITI drawn from a truncated exponential distribution with a minimum of 5.5 seconds, mean of 7, and a maximum of 15.4. There were four unique Gabor cues used in the task that were identical except in the orientation they were presented. The CS+ was pseudo-randomly counterbalanced between-participants to be either 15° or 75° counterclockwise from vertical. The generalization stimuli (GS) are labeled in relation to the orientation of the CS+, such that if the CS+ is 15° then GS1 is 35°, GS2 is 55°, and GS3 is 75°. Each cue took up 5° of visual angle and had a spatial frequency of 3.5 cycles per degree of visual angle.

For each block, the same Gabor cue presentations were randomized, and the same cue could only be presented twice consecutively. After each block, participants used the clicker to rate valence, arousal, and shock expectancy per cue as already described in the procedure. The first block was Habituation, in which no US shocks were given, and each Gabor orientation was presented 8 times for a total of 32 trials. Each subsequent block contained 12 cue presentations for a total of 48 trials. For the first Acquisition block, the first 6 CS+ presentation had a reinforcement rate of 100% which means they were always paired with US ankle shock. The shock was delivered after 2 seconds from the CS+ onset and co-terminated with the cue offset. The first 6 CS+ trials were also boosted such that no more than 2 GS separated CS+ trials. Following the initial CS+ trials, the reinforcement rate was lowered to 50% for the remainder of Acquisition Block 1 as well as for Acquisition Block 2. The last block was Extinction in which the CS+ cue was no longer paired with the US.

### 2.4 | EEG Acquisition and Data Reduction

The EEG was acquired using a 32-channel MR-compatible EEG system (BrainAmp MR;Brain Products). The system uses 31 Ag/AgCl scalp electrodes positioned at the standard 10-20 positions, with the reference at the FCz location and an online bandpass filter from 0.1 to 250 Hz. The 32^nd^ channel records ECG from a channel positioned on the back, slightly to the left of the spine. Processing was accomplished with custom MATLAB scripts using the EEGLAB toolbox and plugins as well as functions from the FreqTag toolbox (Figueira et al., 2022). The data and code used for processing, models, analyses and figures can be found on this project’s OSF page (https://osf.io/hfbjp/).

Two main artifacts effect EEG data in an fMRI scanner: gradient artifacts from the functional scanning and cardioballistic artifacts accentuated by the static magnetic field (Debener et al., 2006). To correct for these artifacts, a mix of specialized equipment and established processing procedures were implemented. To be able to correct for these artifacts, it is necessary to ensure that there are accurate markers for gradient switching and heart beats, such that these repeated sources of error can be subtracted out (Allen et al., 2000; Niazy et al., 2006). The brain products EEG system uses specialized hardware to receive TTL pulses for each fMRI repetition time (TR) which corresponds to each collected volume. These TRs are passed to a synchronization box, to align the MRI and EEG systems clocks, and are then written into the EEG file as markers in the acquired data. The distance between TR pulses was confirmed to be accurate to the 4^th^ decimal place. The data was acquired at 5 KHz, as higher sample rates help in correcting for artifacts. The gradient artifact correction was applied using the fmrib_fastr function described in Niazy et al. (2006) implemented from a EEGlab toolbox script. The function is designed to align volumes, subtract a moving average, and then to subtract principal components of residual artifacts. The PCA removal was implemented but led to no differences in the overall pattern of effects for all three models and thus was omitted from the final pipeline used for the results reported below. Because no PCAs were removed, gradient artifacts were handled by the standard subtraction method as described in Allen et al. (2000). The data were then down-sampled to 500 Hz and the Niazy et al., (2006) algorithm was used to find the R peaks from in the ECG signal, followed by applying the fmrib_pas function which identifies PCA components in the EEG that correspond to cardio-ballistic artifacts. Three principal components were removed to correct for the cardio-ballistic artifact. Processing the data without any cardio-ballistic artifact correction led to near-identical results for all three models with an average of 8 more trials retained per participant. The correction was applied in the present analysis pipeline because it is standard practice (Petro et al., 2017). The data were then filtered with a Hamming windowed sinc FIR with a high-pass of .05 Hz and low pass of 40 Hz (3-dB points), implemented with the pop_eegfiltnew function from EEGlab.

The data were then inspected by two members of the research team independently by scrolling through the recordings and considering the ssVEP spectrum for each trial. Bad channels were agreed upon based on agreement between the research team and are listed per participant in the preprocessing script on OSF (a002EEG_prepro.m; https://osf.io/hfbjp/). No participants had Oz as a bad channel which was the only sensor analyzed. The preprocessing script then performed an ICA using the SOBI algorithm from the pop_runica function in EEGlab (Tran et al., 2009). Removing the ICA components related to eye-blinks led to no differences across the models, so ultimate no ICA components were removed from the final data set. Missing channels were interpolated via a spherical spline using the pop_interp function. The data was then re-referenced from the FCz location to an average reference. The pop_clean_rawdata function was then used to identify artifact contaminated sections of the data. The channel rejection options were disabled. Additionally, no high-pass filtering was applied. Artifactual bursts were identified using a threshold of 20 standard deviations and those sections were trimmed from the data set. The Euclidean distance was used to identify windows in which 25% of the window were affected. In a final step, the data were transformed to a current source density (CSD) representation, implemented via the current_source_density function from the ERPLAB toolbox (Lopez-Calderon & Luck, 2014).

Retained single trials were extracted from each participant via the pop_epoch function. Trials were segmented from cue onset to offset (2000 ms duration). The oscillatory power for the 15 Hz ssVEP driving frequency was quantified via by means of a Discrete Fourier Transform (DFT) implemented from the freqtag_FFT3d function from the FreqTag toolbox (Figueira et al., 2022), applied to the entire 2000 ms segment for each trial, with no windowing applied, leading to a spectral frequency resolution of .5 Hz. This was done for the preprocessed data and was compared against an alternative version of the pipeline in which a sliding window analysis was used to further isolate the ssVEP frequencies (Morgan et al., 1996; Wieser et al., 2016; Figueira et al. 2022). Ultimately, the DFT was used for simplicity because the sliding window analysis led to near equivalent results across the three models. Finally, the mean amplitude for the 15 Hz frequency was extracted for each retained trial from the Oz sensor and was used as the dependent measure for all Bayesian multilevel models (referred to as *Amplitude* in the equations).

### 2.5 | Bayesian models and analyses

The necessary elements for a Bayesian model of empirical data include specifying 1) a model for how the data is distributed (sampling model that captures likelihoods) and 2) priors for all unknown parameters. In line with best practices (Gelman et al., 2014; McElreath et al., 2020) the models were built iteratively, and priors were selected to be uninformative but on a realistic scale for the data (Z-scored ssVEPs). Visualizations and rationale for the uninformative priors can be found in the supplemental materials. Three models were specified and estimated in the Stan statistical programming language (Stan Development Team, 2024) using a variant of the Hamiltonian Monte Carlo sampling method (Duane et al., 1987; Hoffman & Gelman, 2014). A total of 80,000 samples were taken to estimate the posteriors across 8 Markov Chains. For all models, the chains converged with the *R-hat* metric being below 1.05 for all estimated parameters. Five additional models were fit and can be found in the supplemental materials but are not presented in the main text for simplicity and interpretability. Three of the unreported models used slightly different parametrizations of learning ratings leading to lower cross-validation accuracy, but no notable changes in model interpretations. The other two models not reported in the main text used individual participant ratings of arousal and US expectancy per cue and block to predict ssVEP amplitude. There was a marginal relationship between arousal ratings, but poorer cross-validation accuracy compared to the three presented models. The code and discussion of these five models can be found on the OSF project accompanying this manuscript at https://osf.io/hfbjp/.

There are several elements that were held constant for interpretability between the models. The first is that all models were fit to observed and the estimated missing trials. So an array was formed with missing and observed ssVEP amplitudes.

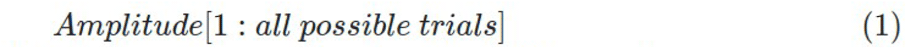

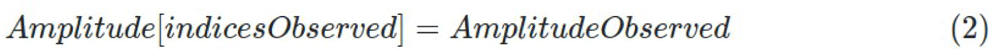

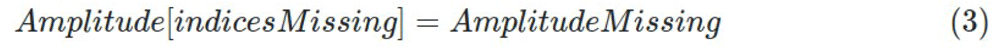

The error for each prediction (*μ*) was specified as Gaussian and was allowed to vary by participant. So were *i* loops through all possible observations and *participant* is an array of integers specifying the current participant:

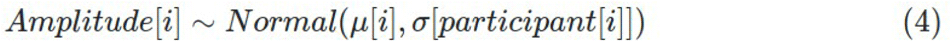

The parameter for the error per participant (*σ*) was given a multilevel prior to regularize it by the average (*σAverage*). Error posteriors closer to zero suggest better predictions per trial. Because of the limited sample size, the regularizing multilevel prior was specified as a Student-T distribution as opposed to a normal distribution. The prior for *σAverage* was centered at the initial Z-scored error of 1.

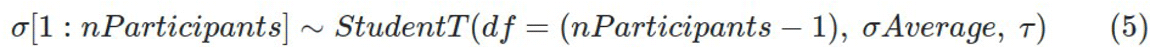

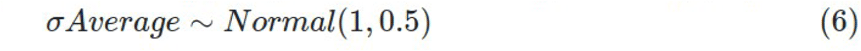

Deviation parameters such as *τ* could be very close to zero if the error per trial between participants (*σ*) is similar between participants. This can make it difficult to sample *τ* with the Stan Hamilitonian Monte Carlo method. For this reason, many of the deviation parameters were specified as an unbounded parameter which were then exponentiated to get back to correct positive only scale required. So as shown, only the *Raw* variant received a prior and were sampled directly before being transformed. This allows for posteriors to be sampled more effectively because they are flatter even if they become very close to zero as the model is fit. As mentioned, visualization and rationale for these priors can be found in the supplemental materials.

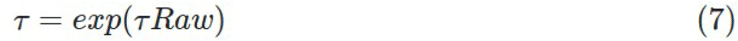

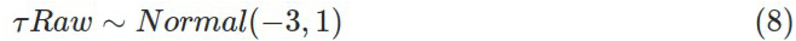

#### 2.5.1 | Model 1: only adaptation over trials with no effect of cue

Prior conditioning studies and other ssVEP studies have observed that the amplitude of visuocortical responses may decrease over the course of the experimental session, thought to reflect several neurophysiological and behavioral factors, including visuocortical adaptation, increasing spontaneous alpha oscillations, and participant fatigue (Norcia et al., 2015; Liu et al., 2012). Thus, we first modeled an adaptation gradient of the ssVEP amplitude over the course the study that could differ between participants. This was modeled as a linear effect across trials for each prediction *μ* for all models.

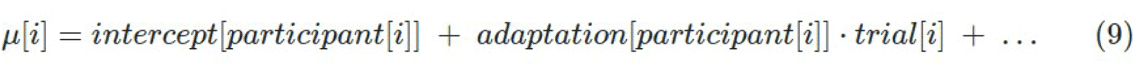

The adaptation intercept and slope was the only predictors used for Model 1, thus this model acts as a Null model for comparative purposes. Because the data is Z-scored within participant, there must be a negative correlation between the *intercept* and *adaptation* slope. Thus, the two parameters were sampled from a multivariate Student-T distribution where *ρ* is converted to be between 0 to -1 with an -inverseLogit function.

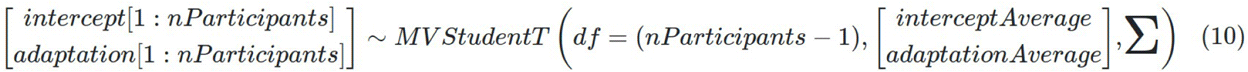

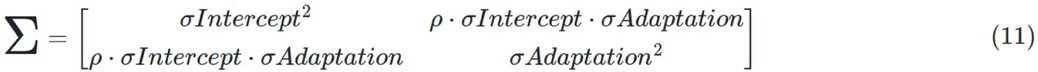

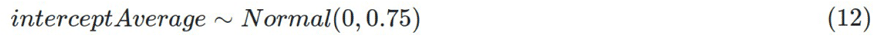

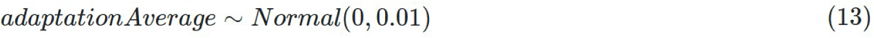

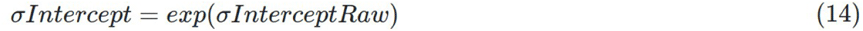

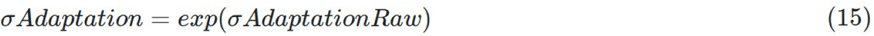

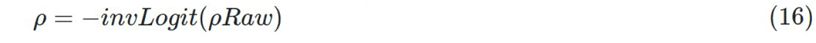

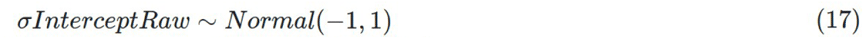

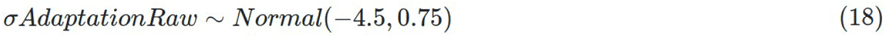

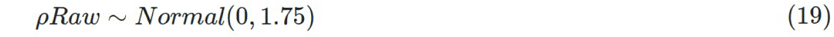

For the same reasons as *τ*, *σIntercept* and *σAdaptation* were estimated via exponentiation of sampled unbounded parameters. The correlation *ρ* was also first specified as unbounded as it is expected the correlation could be very close to -1. The prior for *ρRaw* takes a roughly uniform shape after the -inverseLogit transformation.

#### 2.5.2 | Model 2: mean of cue per block

The second model predicts each trial’s ssVEP as a combination of the adaptation slope with an effect of each cue (CS+, GS1, GS2, GS3) per block (4 total: 1 habituation, 2 Acquisition, 1 Extinction). The effect of each cue per block was specified as a 4×4 array (*βCue*) and was regularized via a Student-T distribution and the deviation parameter *σCue*. The mean for all cues was set to zero to allow it to vary from the adaptation regression line per participant.

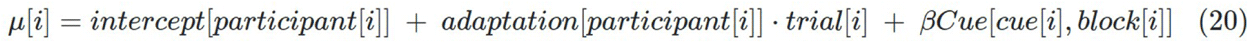

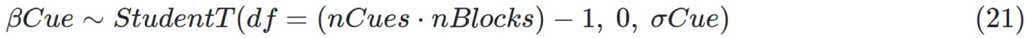

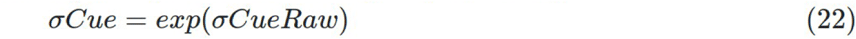

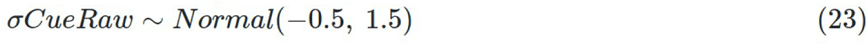

#### 2.5.3 | Model 3: multilevel Rescorla-Wagner inspired Associate Strength

A goal of the present study was to apply models of associative learning to neurophysiological data. The Rescorla-Wagner model is one of the most well-established in which the change in associative strength (*ΔCS+Strength*) is modeled as the difference in current associative strength (*CS+Strength*) from a binary variable specifying if the US was paired or not (1 = US or 0 = no US) scaled by a *LearningRate* parameter (Rescorla & Wagner, 1972). Both *LearningRate* and *CS+Strength* are constrained to be between 0 and 1.

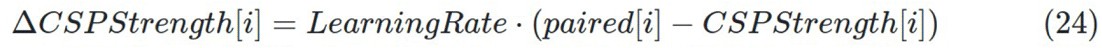

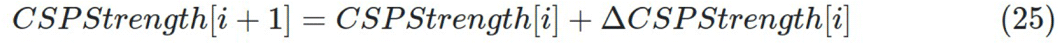

This was augmented to predict ssVEP amplitudes per trial by scaling the *CS+Strength* per cue (4-d array *βScaling*) and allowing the *LearningRate* to differ between participants. It was found that cross-validation was highest when separate learning rates were specified for CS+ trials that were paired and unpaired with the US (*LearningPaired* & *LearningUnpaired*). Additionally, using the invLogit transformation of learning rates also led to higher cross-validation. Thus, the final model structure was:

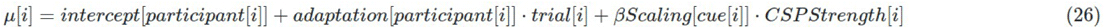

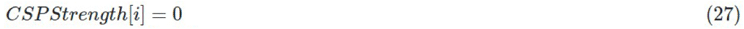

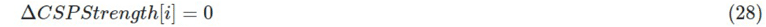

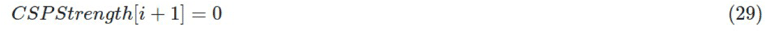

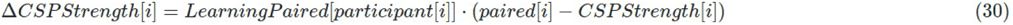

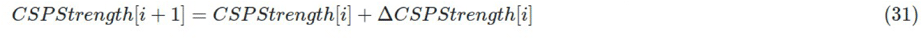

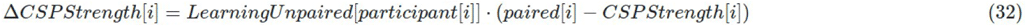

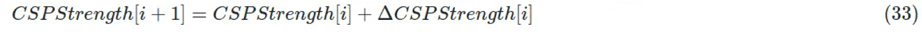

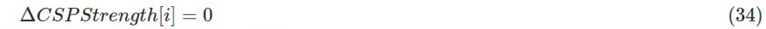

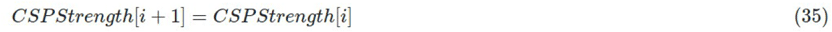

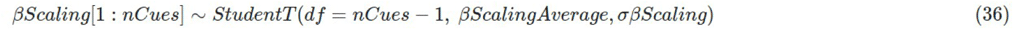

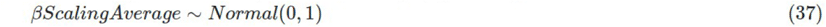

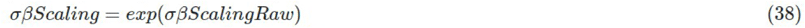

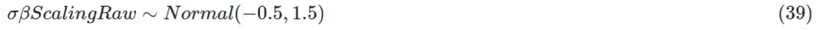

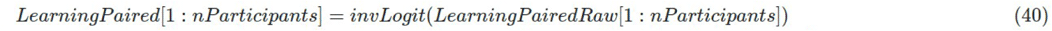

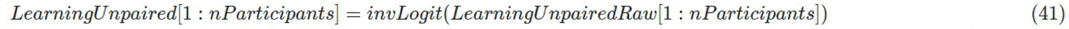

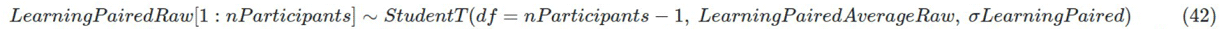

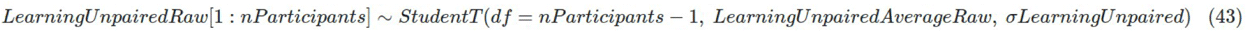

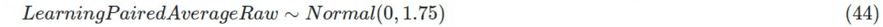

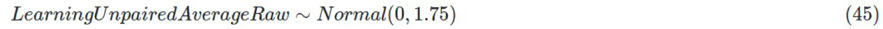

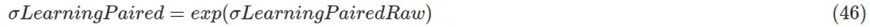

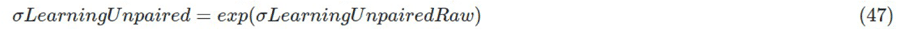

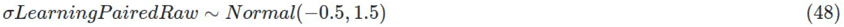

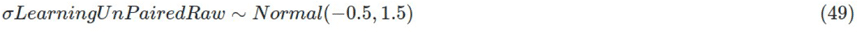

#### 2.5.4 | Model comparison via PSIS-LOO cross-validation

Models with many parameters can overfit to the observed data leading to interpretations that would not generalize to new samples. Cross-validation is an established method for addressing this issue, in which accuracy is measured on data held out of the model fitting. For Bayesian models, the leave one out (LOO) cross-validation accuracy can be estimated with Pareto-smoothed Importance Sampling (PSIS-LOO; Vehtari et al., 2024). This measure has been shown to provide equivalent information to ordinary LOO and k-fold cross-validation or information criteria statistics (Vehtari et al., 2017). Additionally, the models do not need to be iteratively refit and diagnostic Pareto-*k* weights indicate if the cross-validation measure is valid or is being skewed via outliers. Cross-validation accuracy was only measured on observed samples and not the *AmplitudeMissing* parameters used in model estimation. Out of the 3,560 observed ssVEPs, 7 had Pareto-*k* weights above 0.7 indicating outliers for Model 3. However, the estimated effective samples was still high (2,121) and excluding these observations have no effect on overall interpretations.

The PSIS-LOO method converts the log-likelihood per observation to an expected log-likelihood of if the data was held out of the model fitting. Models are compared via the sum of expected log likelihoods known as the expected log-likelihood predictive density (ELPD). Because log-likelihoods are on a scale from -infinity (maximum unlikely) to zero (perfect prediction with no uncertainty), a model with a more positive ELPD indicates better cross-validation accuracy over all observations. Models were compared based on their overall differences in ELPD via built in Stan functions. The standard error of each sum or difference of sums was found with the standard equation:

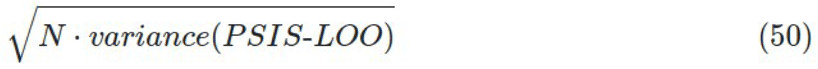

The ELPD values were used to further compare models via stacking weights that indicate what linear combination of the models would lead to best cross-validation accuracy. During this procedure, the number of effective parameters is estimated which can give a sense of how multilevel priors regularize the number of degrees of freedom (Vehtari et al., 2017). ELPD was separated per cue, block, and participant to assess at which points models had better cross-validation accuracy. Finally, the cross-validation accuracy was further interpreted per participant via LOO-adjusted *R^2^* (variance explained) posteriors. This was accomplished via a Dirichlet weight resampling Bayesian Bootstrap procedure adapted from the loo_R2 function from the rstanarm R package (last accessed 12/30/24, https://raw.githubusercontent.com/stan-dev/rstanarm/refs/heads/master/R/bayes_R2.R). After resampling, the *R^2^* is defined as:

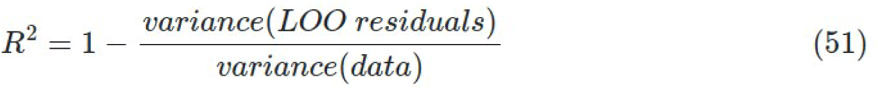

Thus, *R^2^*posterior samples can be negative if the model predictions are poor and produce more variance than in the original data.

#### 2.5.5 | Posterior analyses

Not all estimated parameter posteriors are presented in the main text, but they can be found in the supplemental materials. In line with contemporary statistical recommendations (McElreath, 2020; Wasserstein et al., 2019), representations of statistical uncertainty with the posteriors were depicted or numerically described as opposed to Null hypothesis tests. The learning rate posteriors per participant were plotted as densities between the values their bounds of 0 and 1. To show how learning rates relate to *CS+strength*, the median and 100 posterior draws of this posterior are also depicted. To understand if specific learning patterns were related to poor model fit, the LOO-adjusted *R^2^* posteriors per model were also depicted by participant in the same figure (Figure 3). To understand the effects of cue and US pairing, the posterior densities for *βCue* from Model 2 and *βScaling* from Model 3 are depicted. In this same figure the combined effect of *βScaling* with *CS+Strength* are demonstrated via product of these parameters for three participants (1, 5, and 14) via 100 posterior draws and the full posterior average per cue. Lastly, the missing trial interpolation for Model 3 is illustrated by 100 draws for the missing trials in a subsection of the data fro participants 1 and 14. These interpolated trials are contextualized with lines representing the average posterior for the mean per cue and the surrounding observed data points.

## Results

### 3.1 | Z-scored ssVEP per Participant

The 24 included participants had an average of 148.3 (SD = 22.6) retained trials after preprocessing. The Z-scored mean ssVEP amplitude is depicted in Figure 1.

**Figure 1.**
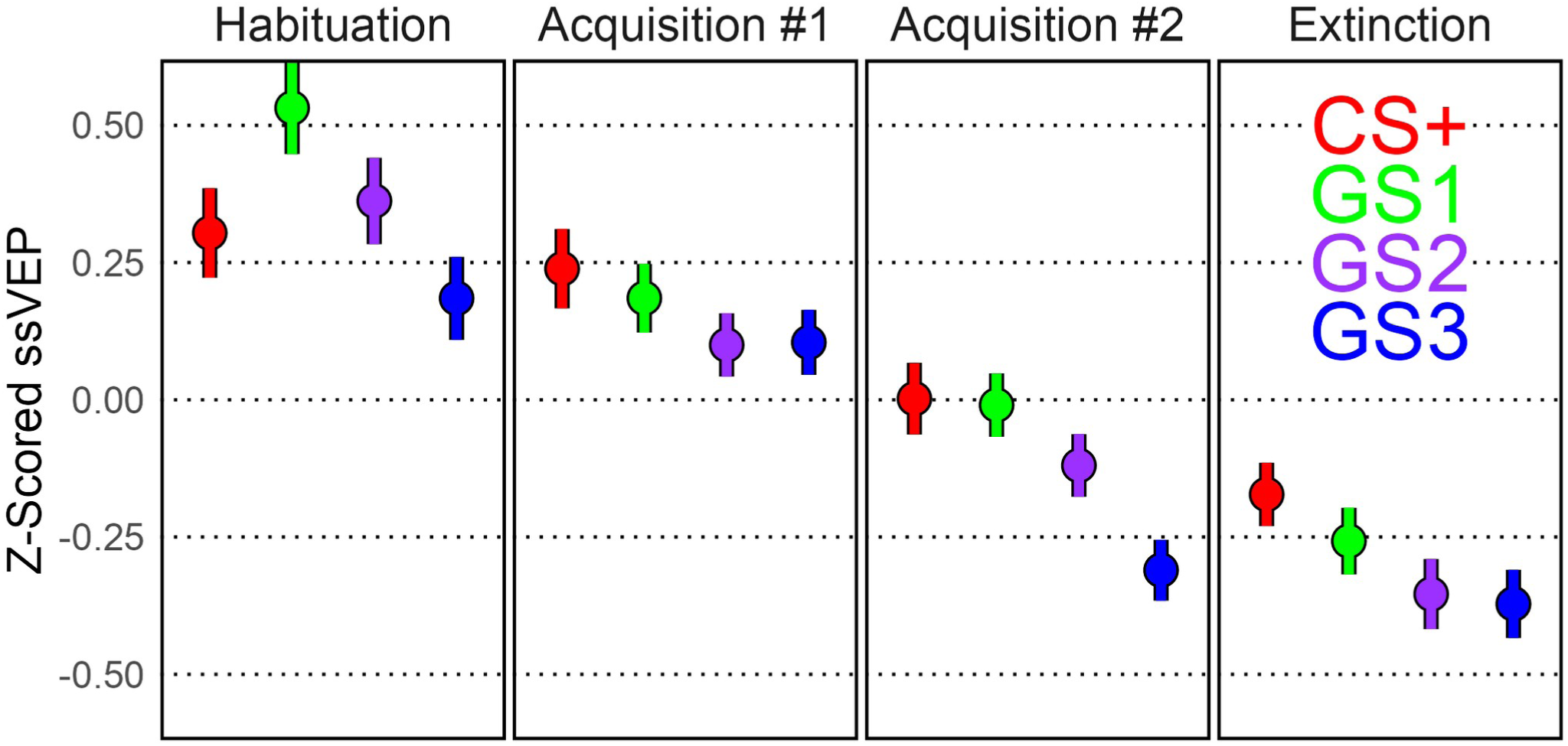
Z-scored ssVEP mean and standard error per block and cue. The CS+ was paired with a noxious electric ankle shock (US) starting in the first Acquisition block at a reinforcement rate of 100% for the first 6 trials and down to 50% after. The generalization stimuli (GS) were never paired with the US. The GS1 was closest in orientation to the CS+, followed by the GS2 and GS3 accordingly.

### 3.2 | Model Comparison via Cross-Validation Accuracy

The PSIS-LOO method found the Learning Model (Model 3) had the best cross-validation accuracy compared to Model 1 and 2, as well as all other supplemental models (Table 1). This is further shown in the model weights, in which the most cross-validation accuracy would be to base 95% of the predict on the Model 3. When Model 3 is excluded, cue by block model (Model 2) outperforms the null model (Model 1) with a similar difference in model weights.

**Table 1.**
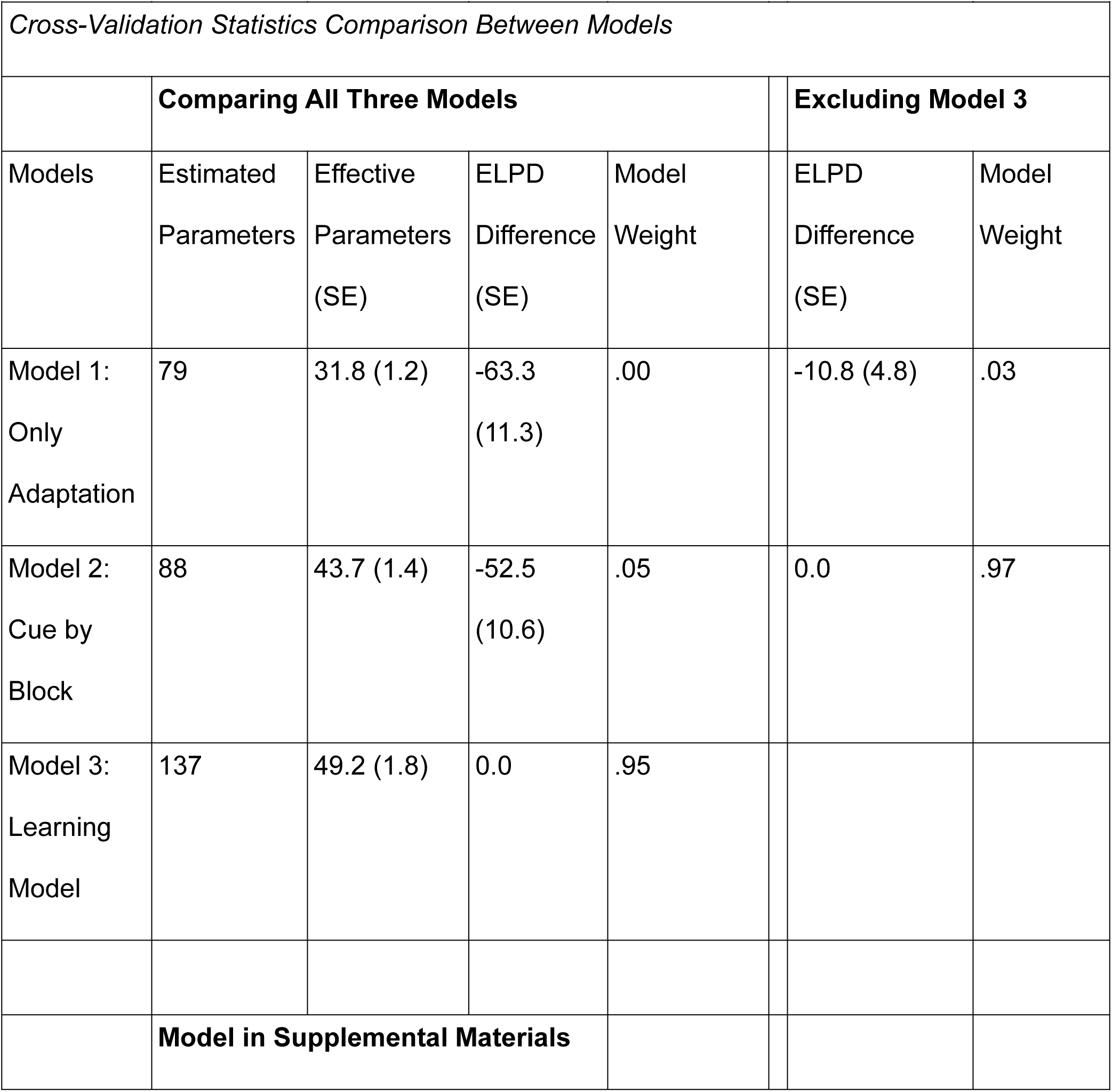
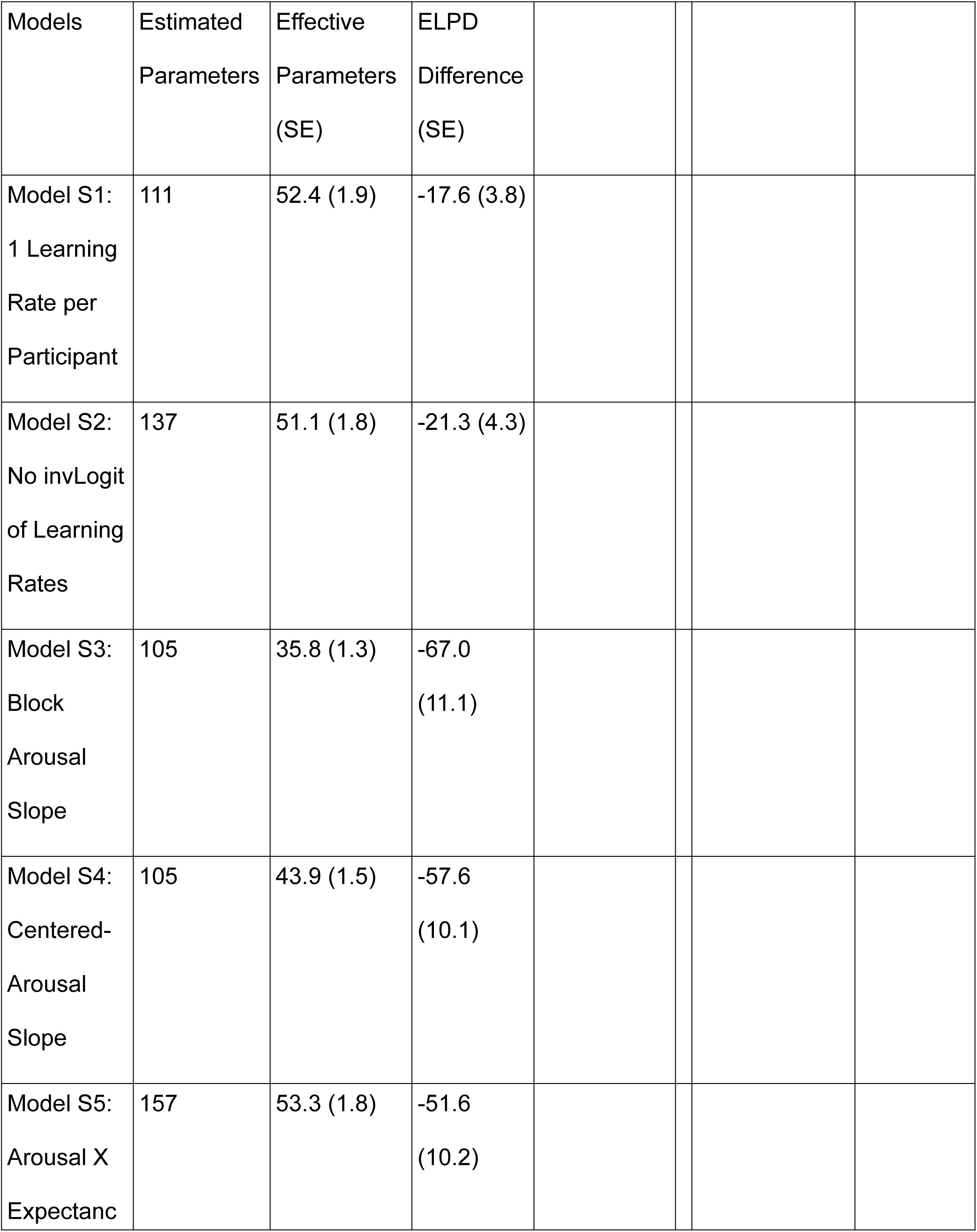
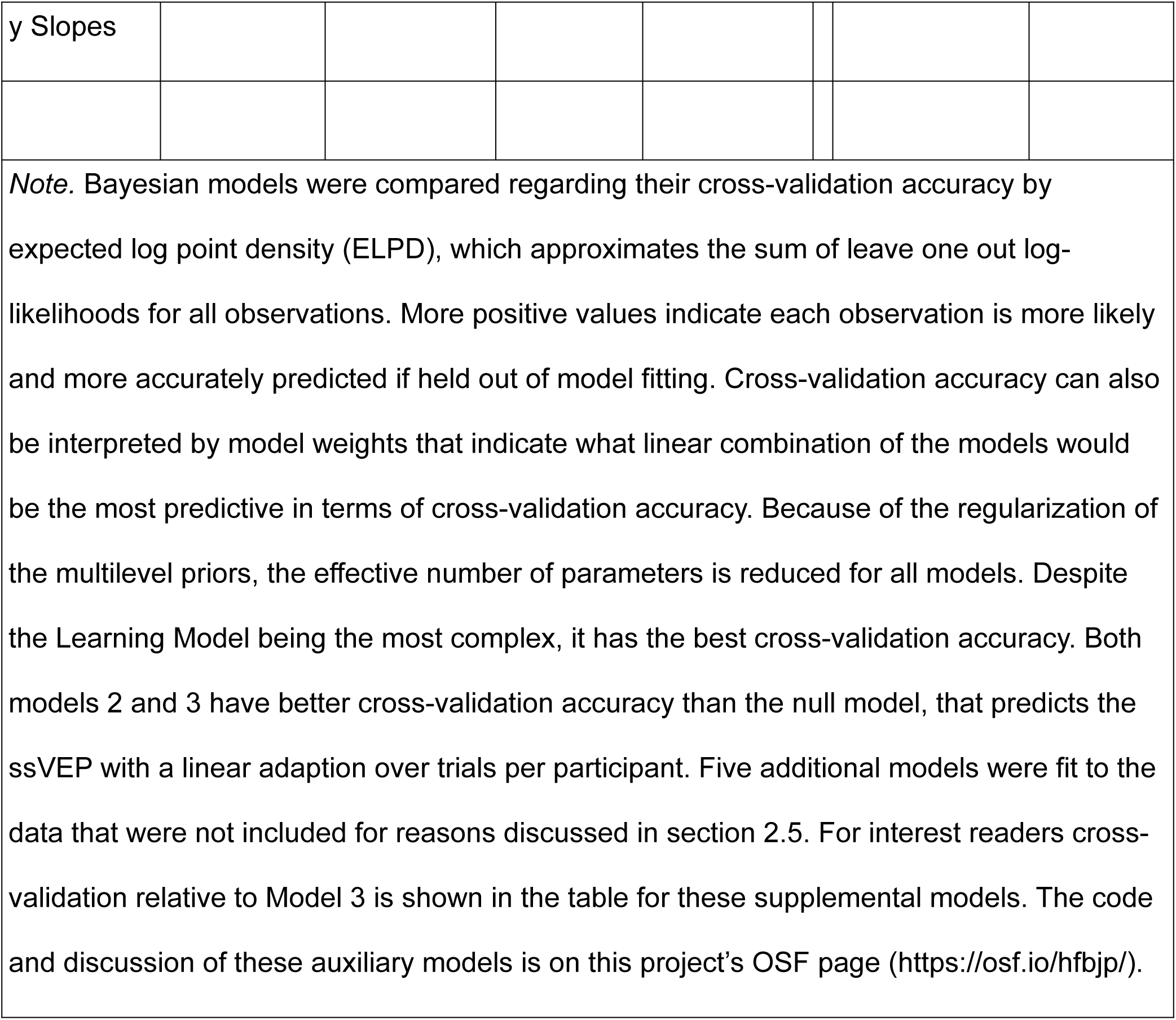

|  |  |  |  |  |  |  |
| --- | --- | --- | --- | --- | --- | --- |
| **Table 1** |  |  |  |  |  |  |
| *Cross-Validation Statistics Comparison Between Models* |  |  |  |  |  |  |
|  | **Comparing All Three Models** |  |  |  | **Excluding Model 3** |  |
| Models | Estimated<br>Parameters | Effective<br>Parameters<br>(SE) | ELPD<br>Difference<br>(SE) | Model<br>Weight | ELPD<br>Difference<br>(SE) | Model<br>Weight |
| Model 1:<br>Only<br>Adaptation | 79 | 31.8 (1.2) | -63.3<br>(11.3) | .00 | -10.8 (4.8) | .03 |
| Model 2:<br>Cue by<br>Block | 88 | 43.7 (1.4) | -52.5<br>(10.6) | .05 | 0.0 | .97 |
| Model 3:<br>Learning<br>Model | 137 | 49.2 (1.8) | 0.0 | .95 |  |  |
|  | **Model in Supplemental Materials** |  |  |  |  |  |

| Models | Estimated<br>Parameters | Effective<br>Parameters<br>(SE) | ELPD<br>Difference<br>(SE) |
| --- | --- | --- | --- |
| Model S1:<br>1 Learning<br>Rate per<br>Participant | 111 | 52.4 (1.9) | -17.6 (3.8) |
| Model S2:<br>No invLogit<br>of Learning<br>Rates | 137 | 51.1 (1.8) | -21.3 (4.3) |
| Model S3:<br>Block<br>Arousal<br>Slope | 105 | 35.8 (1.3) | -67.0<br>(11.1) |
| Model S4:<br>Centered-Arousal<br>Slope | 105 | 43.9 (1.5) | -57.6<br>(10.1) |
| Model S5:<br>Arousal X<br>Expectanc | 157 | 53.3 (1.8) | -51.6<br>(10.2) |

|  |
| --- |
| y Slopes |

*Note.* Bayesian models were compared regarding their cross-validation accuracy by expected log point density (ELPD), which approximates the sum of leave one out log-likelihoods for all observations. More positive values indicate each observation is more likely and more accurately predicted if held out of model fitting. Cross-validation accuracy can also be interpreted by model weights that indicate what linear combination of the models would be the most predictive in terms of cross-validation accuracy. Because of the regularization of the multilevel priors, the effective number of parameters is reduced for all models. Despite the Learning Model being the most complex, it has the best cross-validation accuracy. Both models 2 and 3 have better cross-validation accuracy than the null model, that predicts the ssVEP with a linear adaption over trials per participant. Five additional models were fit to the data that were not included for reasons discussed in section 2.5. For interest readers cross-validation relative to Model 3 is shown in the table for these supplemental models. The code and discussion of these auxiliary models is on this project’s OSF page (https://osf.io/hfbjp/).

A breakdown of ELPD differences by cue, block, and participant illustrates at which points Model 3 had better cross-validation accuracy than Model 2 (Figure 2). A majority of the cues per block and participants were better predicted by Model 3. However, the bulk of Model 3’s ELPD improvement was because of the CS+ prediction in blocks following habituation and for a few specific participants.

**Figure 2.**
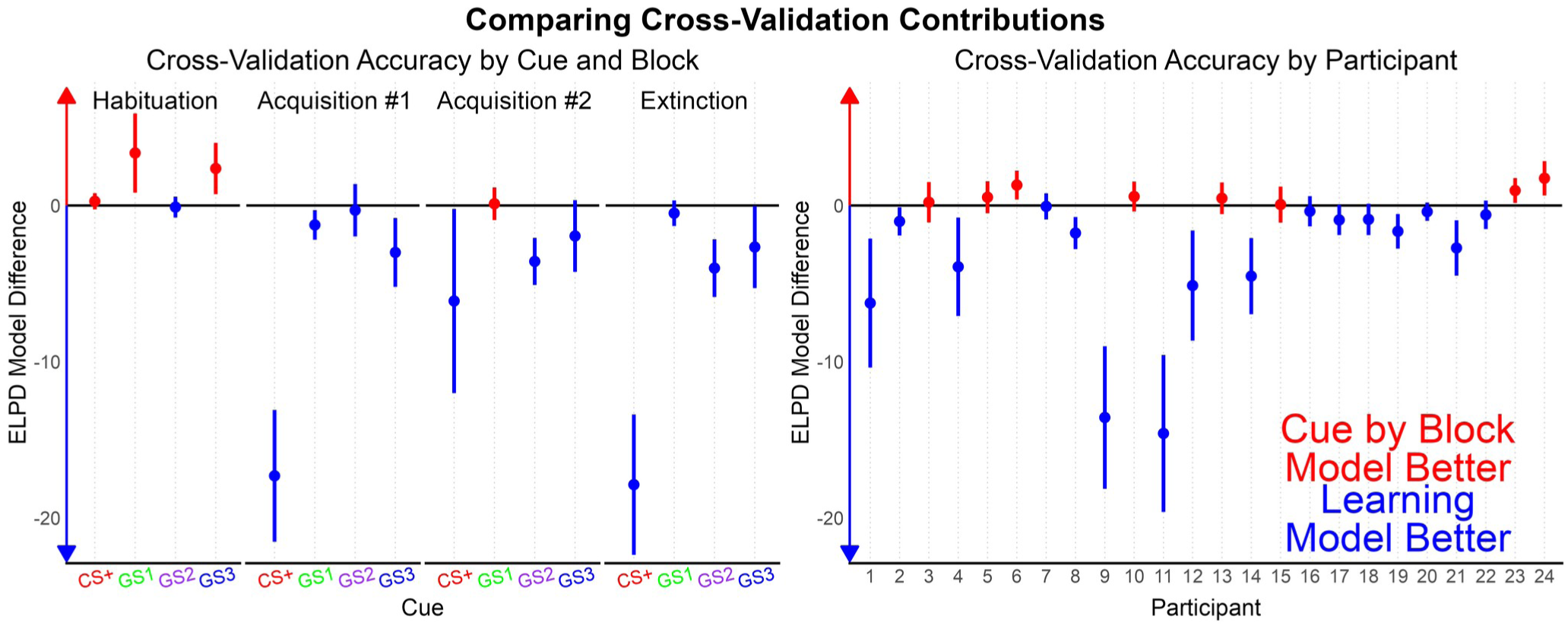
The cross-validation measure ELPD broken down by cue, block and by participant. ELPD is the sum of log-likelihoods per observed data point if that data point were held out of model fitting. Visualizing the difference in ELPDs between models across relevant experimental conditions is recommended for understanding why models are more accurate and how they could possibly be improved. By subtracting the learning model (Model 3) from the cue by block model (Model 2), it becomes clear that the learning model is primarily more accurate for the CS+ cue after the habituation block and for specific participants.

### 3.3 | Posterior Analysis

The downward trend in ssVEP over trials seen in Figure 1 was confirmed via the adaptation intercept and posteriors not overlapping with zero in each of the three models: Model 1: intercept = 0.44 (SD = 0.07), slope = -0.005 (SD = 0.0007); Model 2: intercept = 0.52 (SD = 0.09), slope = -0.006 (SD = 0.001); Model 3: intercept = 0.44 (SD = 0.07), slope = -0.005 (SD = 0.0008). Because the adaptation was similar between models, it is not further described in the main text. However, a figure of the adaptation (Figure S1) and full posteriors per participant can be found in the supplemental materials.

Model 3 posteriors relevant to associative strength learning (Figure 3) suggest different learning patterns between-participants. The average learning rate of CS+ pairings with the US (*LearningPaired*) had a median of .22 (95% credibility interval: [.01, .77]) whereas the *LearningUnpaired* for unpaired CS+ trials was .33 [.02, .87]. The posterior for learning rate deviations between participants increased from a prior of Normal(−0.5,1.5) to a mean of 2.58 (roughly normally distributed; SD = 0.87) for *σLearningPairedRaw*, and to 2.09 (SD = 0.95) for *σLearningUnpairedRaw*. This suggests that there was evidence that the learning rates between participants differed more than the chosen prior. While *CS+Strength* is not an estimated parameter, it too has a posterior dependent on draws from the two learning rates per participant. At trial 37, the first Acquisition block begins in which the first 6 CS+ trials are paired with a US (100% reinforcement rate). After these first 6 boosted trials the CS+ is only paired with a US 50% of the time. The majority of participants have *LearningPaired* posteriors close to 0 with the *LearningUnpaired* evenly distributed at 0 and 1. These specific participant have *CS+Strength* posteriors that differ very little with each CS+ pairing. Five of the participants (1, 4, 9, 11, and 12) have *LearningPaired* rates close to 1 and *LearningUnpaired* close to 0, leading to *CS+Strength* close to 1 throughout the last three experiment blocks.

**Figure 3.**
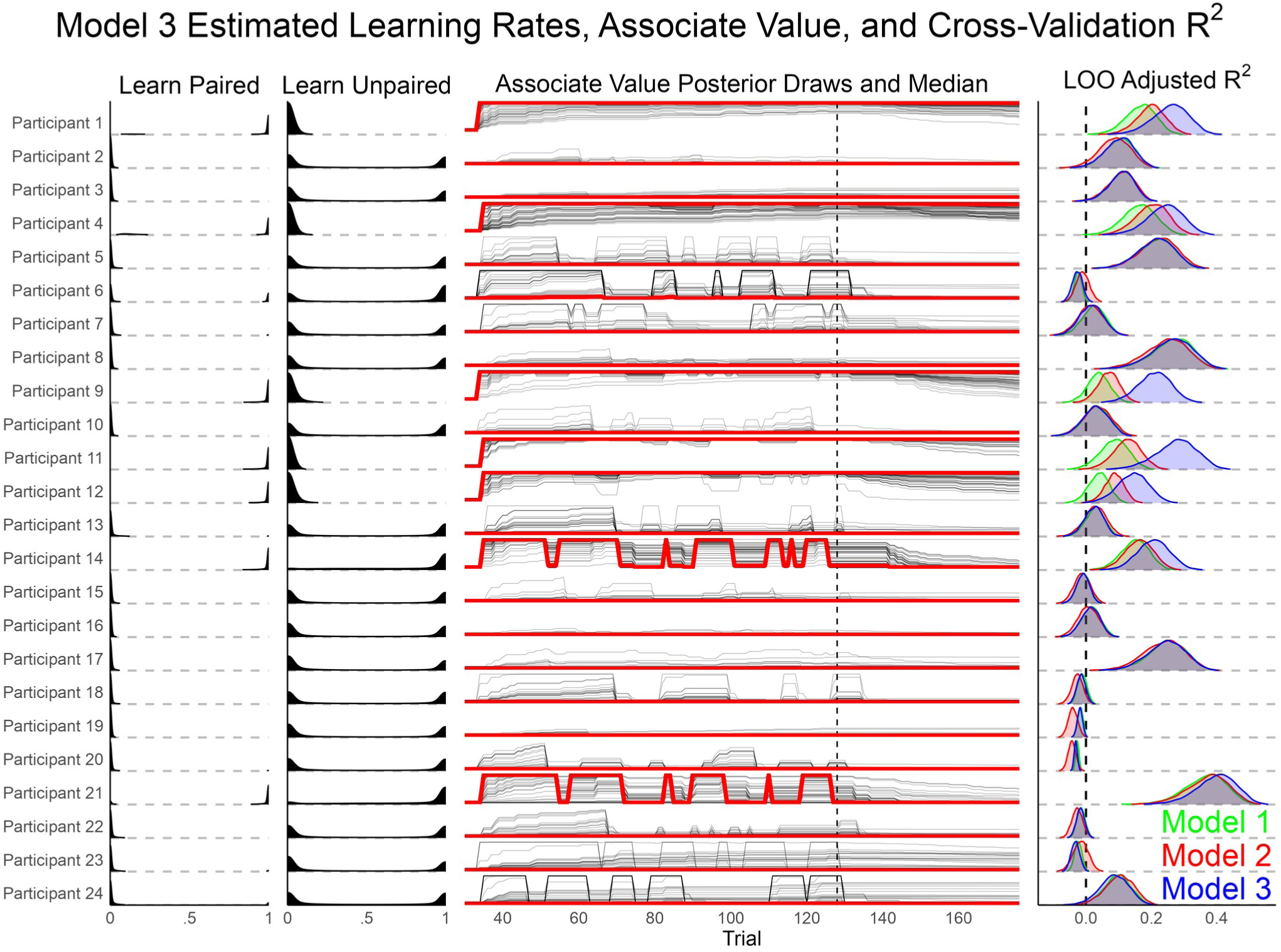
A visualization of learning rates and estimated associate value per participant. From left to right, the *LearningPaired* parameter specifies how quickly associative strength between the CS+ and US changes with each US pairing. The *LearningUnpaired* parameter indicates decreases in associate strength when the CS+ is presented without a US. The effects of these learning rates on associate strength is depicted in the third panel over trials, excluding the habituation trials where *CS+Strength* was zero. Black thin lines are 100 random posterior draws of associate strength, the red line is the median posterior out of all samples, and the vertical dashed line is the beginning of the Extinction phase. For comparison purposes, the cross-validation LOO-*R^2^* is presented for each participant between all three models. Overall, the model predicts a variety of learning patterns across participants with an increase in cross-validation accuracy for most participants that were also predicted well by Models 1 and 2.

Lastly, two participants (14, 21) have a *LearningPaired* posterior close to 1, but a *LearningUnpaired* posterior also close to 1. This led to large shifts in the *CS+Strength* median with each CS+ paired and unpaired trial. Also in Figure 3, the *R^2^* cross-validation posteriors are visualized between the three models to contextualize each participant. A majority of participants that had little change in CS+Strength were not fit well by any of the 3 models with LOO-*R^2^* at or below zero. For participants with LOO-*R^2^* above zero, Model 3 match or exceed the other models in cross-validation variance explained. The overall median and 95% credibility interval LOO-R2 per model was: Model 1 = .09 [.07, .11], Model 2 = .09 [.07, .11], Model 3 = .12 [.10, .14]. A posterior contrast suggest Model 3 explains .031 [.004, .058] more single-trial cross-validation variance than Model 1, and .026 [-.001, .053] than Model 2.

Finally, the parameters most directly tied to differences in ssVEP cue predictions were plotted in Figure 4. *βCue* from Model 2 depicts the average uncertainty per cue per block. Alternatively, *βScaling* from Model 3 is the predicted change in cue when *CS+Strength* is at the maximum value of 1. To illustrate the complete Model 3 cue prediction, the average and 100 posterior draws for each cue prediction are depicted for three participants (Figure 4).

**Figure 4.**
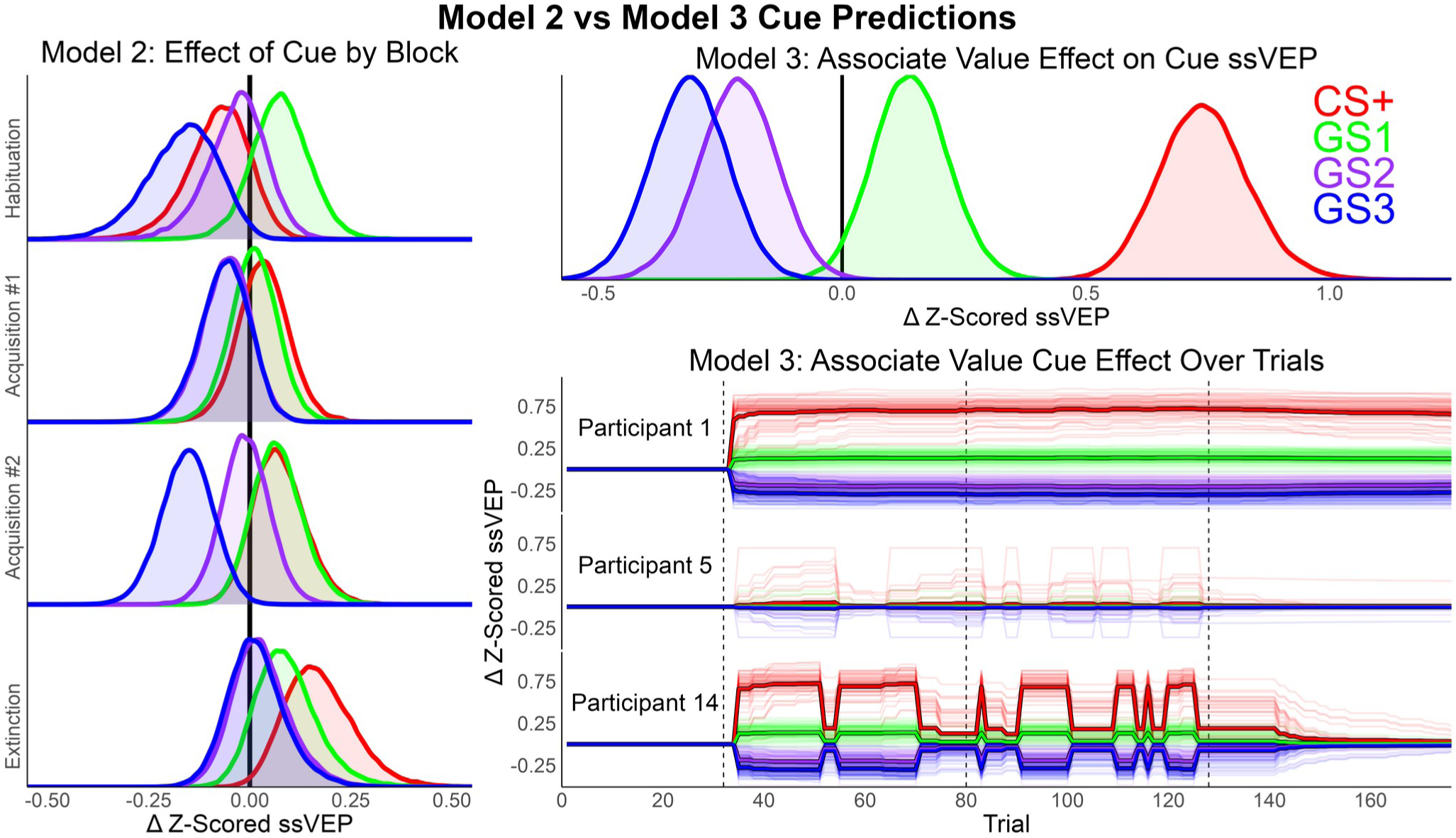
Model 2 and 3 cue relevant posteriors controlling for adaptation. Left: Posteriors for *βCue* from Model 2 represent the probability of change for the ssVEP per cue by block. Top right: The posteriors of *βScaling* from Model 3 estimate the effect on ssVEP per cue when CS+ associate strength is 1. Bottom right: The predicted effect per trial and cue (*βScaling*[1:*nCues*] ⋅ *CS+Strength*[i]) for participants 1, 5, and 14. These participants were selected because they have different patterns of learning, while still having positive LOO-*R^2^*posteriors. Thin lines are 100 random posterior draws, the thicker line is the average of all samples, and vertical dashed lines indicate transition between blocks (Habituation, Acquisition #1, Acquisition #2, and Extinction). In Model 3, the effect of cue conditioning is accentuated because it can differ over trials and between participants based on *CS+Strength* lowering statistical uncertainty.

Lastly the models were used to recreate the original data. The grand mean and its model recreations are depicted in Figure 5. The correlation between observation and model predictions all had *p*-values less than .001: Model 1 *r* = .32, *t* = 20.0; Model 2 *r* = .33, *t* = 21.2; Model 3 *r* = .37, *t* = 23.8.

**Figure 5.**
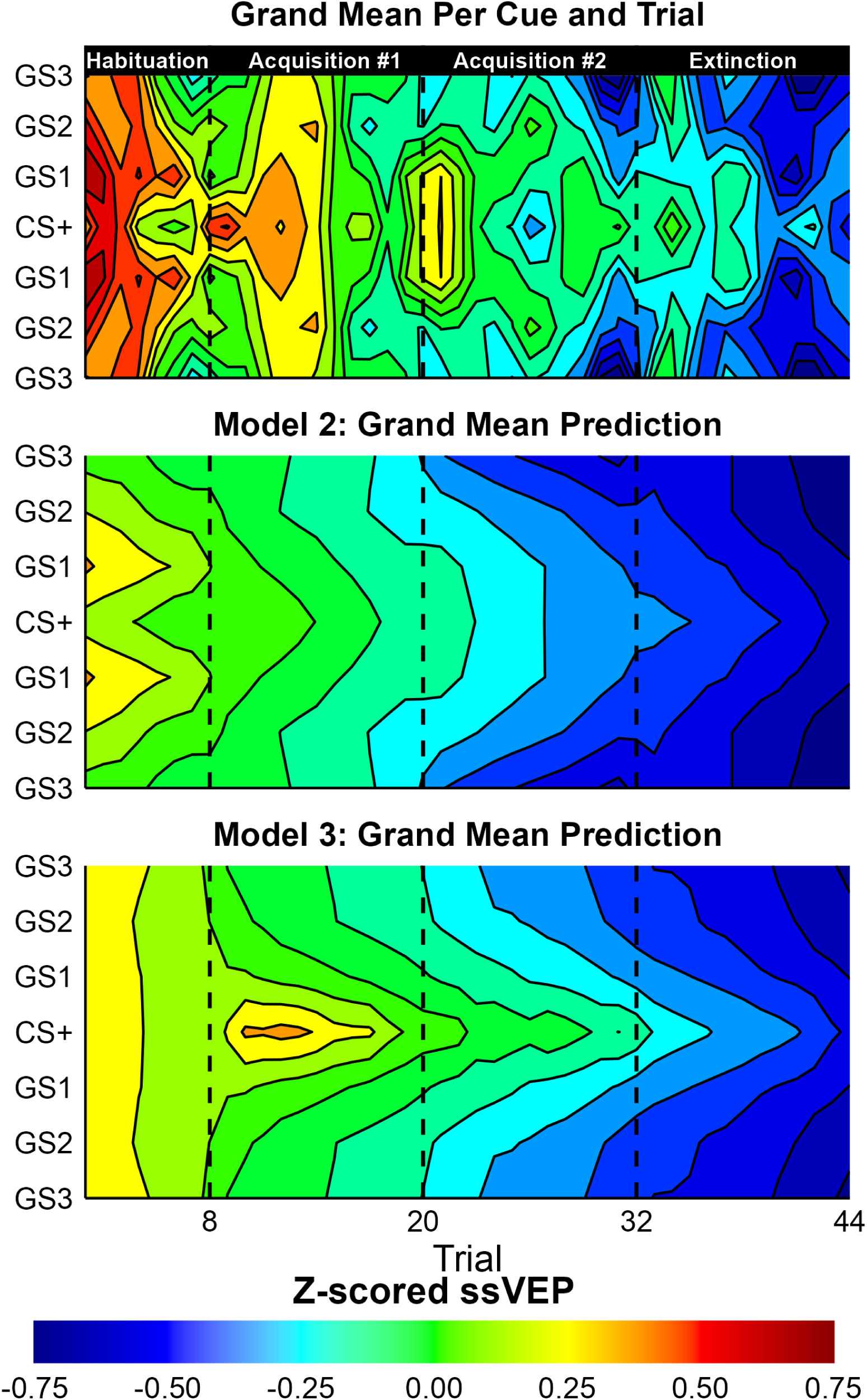
A visualization of the grand mean Z-scored ssVEP observed, and predicted by Model 2 and 3. The contour plot depicts the average of all trials per cue per participant. The grand means per trial were then smoothed with a moving average of two surrounding trials per cue.

## Discussion

The present study used Bayesian multilevel models to quantify the effect of differential conditioning on cue-evoked ssVEP amplitudes. The overarching goal was to establish that a multilevel learning model can accurately recover aversive generalization by overcoming single-trial noise, inter-individual differences, and missing trials. By detailing how models are designed, compared through cross-validation, and analyzed; this project serves as the theoretical basis for future models that will be more ambitious and informative.

Visible in the cue means (Figure 1), there was an overall decrease in ssVEP amplitude over trials, with greater responses to the CS+ and GS1 cues beginning in the Acquisition phase when the CS+ began being paired with the noxious US ankle shock. Three Bayesian Multilevel models were fit to the data with different predictions per trial: 1) a Null model that only accounted for the ssVEP adaptation (decrease) as a multilevel factor by participant, 2) an additional prediction for each cue per block, and 3) Rescorla-Wagner inspired learning model of associate strength that also controlled for adaptation over trials. In line with the suggested best practices for Markov Chain Bayesian models (Gelman et al., 2021; McElreath, 2020), model comparison was performed with the PSIS-LOO cross-validation method that approximates the log-likelihood of each observation when held out of model fitting (Vehtari et al., 2017; 2024). While the Learning Model was the most complex (137 estimated parameters), it had the best cross-validation accuracy, partly because of the multilevel structure constraining the degrees of freedom to lower the effective parameters to 49.2 (SE = 1.8). As is recommended for PSIS-LOO model comparison (Vehtari et al., 2017), the ELPD per observation were segmented by experimental conditions to understand why Model 3 outperformed Model 2 in cross-validation accuracy. While Model 3 had the best cross-validation accuracy for the majority of experimental conditions and participants, most of its cross-validation advantage is because of 1) predictions for the CS+ cue starting in the Acquisition phase, and 2) better fit for a subset of the participants (Figure 2).

After establishing that Model 3 is predictive and unlikely to be overfitting the data, the posteriors of interest were used to assess experimental questions. First, it was found that there was a reliable average effect of adaptation with the ssVEP decreasing over the study (Supplemental Figure 1). This average adaptation and the adaptation per participant was similar in all three models. Next, the learning rate parameters were visualized per participant (Figure 3). Although learning rates could fall anywhere between 0 and 1, the posteriors concentrated at the minimum and maximum. This resulted in 3 general patterns: 1) participants that had little change in *CS+Strength* (*LearningPaired* ≃ 0), 2) participants that quickly approached max *CS+Strength* with each US-paired trial and did not decrease with unpaired trials (*LearningPaired* ≃ 1; *LearningUnpaired* ≃ 0), and 3) two participants with rapid changes in *CS+Strength* (*LearningPaired* ≃ 1; *LearningUnpaired* ≃ 1). This is shown in Figure 3 by the *CS+Strength* posterior, which is dependent on the learning rates posteriors. A relevant question for interpreting these results per participant is, are there differences in cross-validation variance explained (LOO-*R^2^*) between the participants by model? If the learning Model poorly predicts observations for some participants, then the learning rates may not be interpretable. Thus, the posterior of LOO-*R^2^* is plotted for each participant in Figure 3 as well. For about half of the participants, the LOO-*R^2^* posteriors are at or below zero for each of the three models and Model 3 predicts no change in *CS+Strength*. So, for these participants, where Model 3 does poorly, so do the other simpler models. For the rest of the participants that were well-explained (LOO-*R^2^* > 0), the learning model closely matches or is superior to the simpler models. There are examples of participants that are fit well for each of the three identified learning patterns, and all the participants that have a high *LearningPaired* rate are well-predicted. This suggests that *CS+Strength* dependent changes in ssVEP are justifiably interpretable. So, lastly the predicted effects of conditioning on cue are visualized in Figure 4. The learning model demonstrates that there is more statistical certainty (less overlap) for how conditioning affects each cue, and it predicts a large change for each cue compared to Model 2.

It could be argued that this is a form of “cherry picking” in which we let Model 3 choose who the conditioning effect applies to, so of course it accentuates the effect of conditioning. It is true that in hindsight Model 3 applies the conditioning effect selectively to each participant, but this is not being hidden. In fact, the complete set of posterior parameters details a level of specificity and transparency ideal for scientific inference and actionable future directions. It has been argued that replicable science needs to be open and transparent, while also detailing effect sizes and even how many participants show the effect of interest (Shrout & Rodgers, 2018). Here, it is clearly depicted which participants show the conditioning effect, how it changes over trials, and for which participants cross-validation accuracy differs (ELPD) as well as a meaningful effect size measure (LOO-*R^2^*). Much of this information is lost in Model 2 by holding the effect of cue constant between-participants. As shown in cross-validation metrics, Model 2 is a less accurate depiction of each participant by over-estimating the effect of conditioning for those that do not show the effect and underestimating it for those that do. This makes it more difficult to draw actionable insights for future directions, such as understanding why some participants do not show the effect or which learning models may better fit the underlying data. These analyses are in line with modern statistical recommendations to accurately capture the multilevel structure of the data and to represent statistical uncertainty transparently, moving away from null-hypothesis testing (Wasserstein et al., 2019; Gelman et al., 2024).

Lastly, a benefit of this analysis approach is how missing trials are interpolated. It is common for trials to be lost for reasons such as of movement, eye-blinks, or EEG sensors losing contact with the skin. When additional physiological methods are recorded simultaneously, there will be even more trials missing at least one of the measures. For difficult simultaneous EEG-fMRI recordings, it is common to entirely lose either the EEG or fMRI recording for a participant. So while the overall objective of concurrent ssVEP-fMRI recordings is to utilize ssVEP effects to elucidate fMRI activations, traditional methods would sacrifice statistical power and would need to exclude many participants and/or trials, hampering external validity and efficiency. But as depicted (Figure 6) in this Bayesian framework, missing trials are listed as parameters to be interpolated. The entire model is used to produce a distribution of the missing observation based on the trial’s predicted mean (*μ* [i]) and estimated error per participant (*σ* [participant [i]]). In future models, in which EEG and fMRI trials are specified as coming from a multivariate / multidimensional distribution, a predicted missing trial would still inform an observed fMRI trial and vice versa, but it would be weighted appropriately less than observed trials.

**Figure 6.**
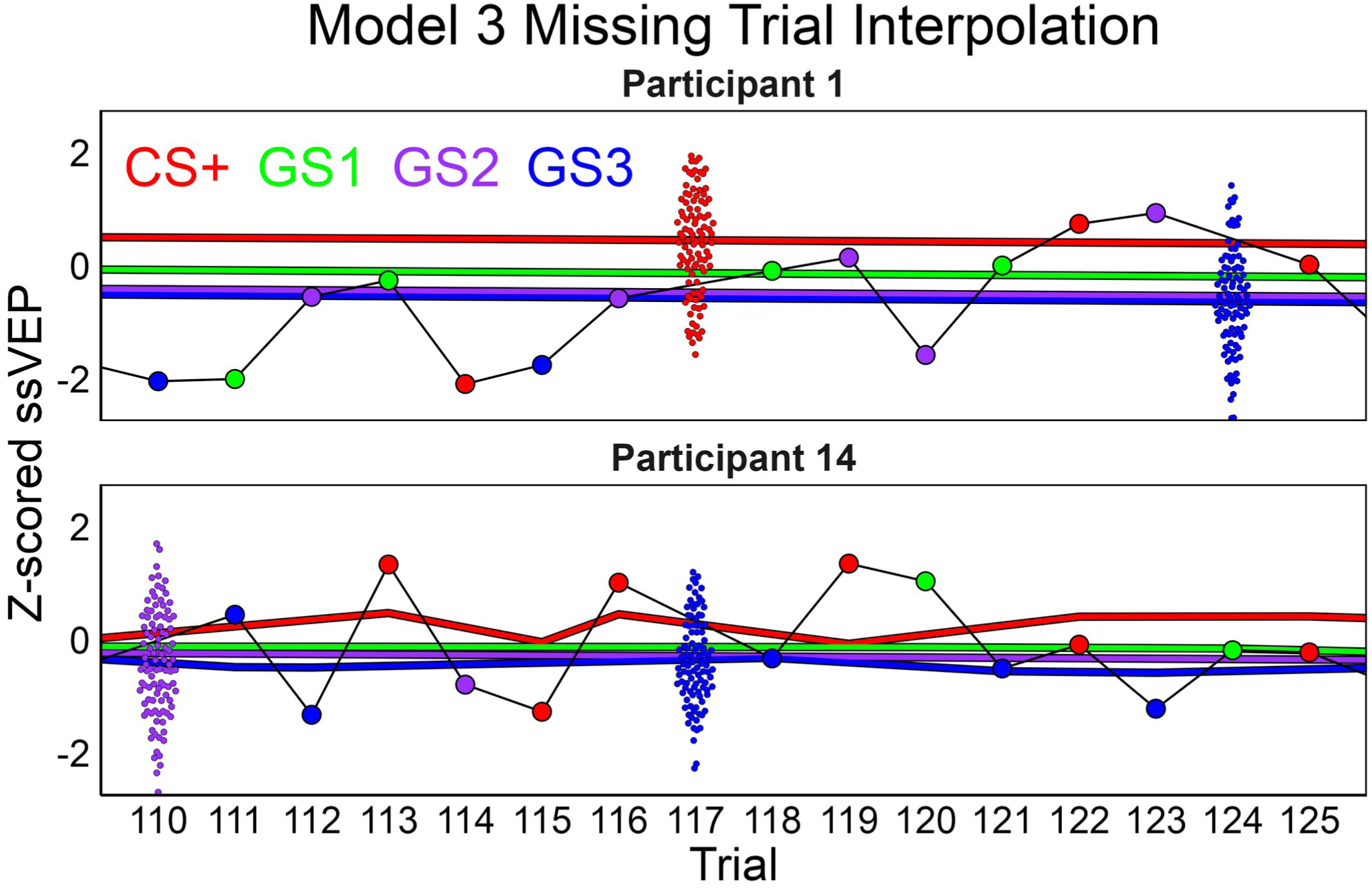
Example of missing trial interpolation for Model 3. Large dots are observed trials, thick lines are the average mean prediction per cue, and swarmed smaller dots are 100 draws for missing trials from the likelihood function. The 100 draws are plotted for clarity, but a complete analysis would use all the draws. In physiological studies, single-trial variability is high and it is common to lose trials for various reasons. This can have debilitating effects on statistical power, particularly for the current project that aims to correlate EEG and fMRI in dynamic learning processes. But in this Bayesian framework, missing trials are listed as parameters and are estimated by draws from the current prediction (*μ* [i]) and estimated single-trial error (*σ* [participant [i]]). This is a justifiable replacement for a missing trial as it act as a compromise, allowing for missing trials to still aid in model fitting while appropriately weighting them less than the actual observed trials.

To efficiently map many dynamic brain processes, the field needs robust single-trial estimates from theory-derived generative models. The benefits of using single-trials for neuroimaging inferences is well-established in the literature (Debner et al., 2007; Ullsperger, 2024), but most of these analyses primarily use single-trials to draw group-level statistical inferences with a limited number of parameters. The analyses demonstrated here not only draw conclusions from single-trials, but provide a justified prediction for a specific trial, cue, and participant without noise (*μ* [i]) or a distribution with noise (*Normal*(*μ* [i], *σ* [participant [i]])). More similar to the current approach are frequentist multilevel and structural equation models that are increasingly being used for neuroimaging data (Volpert-Esmond et al., 2021). These models can be similarly structured and theoretically should reach similar point-estimates when fit by restricted maximum likelihood. However, frequentist multilevel models can be more difficult to fit, and interpreting the parameters tends to involve the interpretation of *p*-values that are often confused and misinterpreted (Wasserstein et al., 2016). Because Bayesian results are representations of the probability of a drawn sample, they are generative models capable of simulating data or interpolating missing trials. These benefits make it a compelling option particularly when obtaining robust estimates of single trials is necessary.

More generally, cutting-edge Bayesian innovations enable the development of explanatory models suitable for scientific inference. While Bayesian statistics and the mathematical principles of multilevel models are historically old and well-established, only in the last 50 years have modern computing and algorithm innovations enabled its widespread use in applied science (Schoot et al., 2021; Gelman & Aki Vehtari, 2021). Even still, algorithms for posterior estimation continue to improve and are packaged with expressive statistical software allowing for quick and flexible implementations of cutting-edge statistical summaries and metrics (Štrumbelj et al., 2024). These iterative improvements in Bayesian software now allow the scaling of theory-driven models to thousands of parameters that have complex non-linear relationships (Gelman et al., 2015; Carpenter et al., 2017). This flexibility, paired with the logic of multilevel modeling, allow researchers to impart more of their domain expertise into their models and a pooling of information from disparate sources while controlling for confounds in detailed ways. Thus, this form of modeling is useful for causal inference in fields such as anthropology in which experiments are not possible (Deffner et al., 2022, 2024; Friederici et al., 2024). Primarily the field of neuroimaging has utilized innovations in machine learning algorithms that maximize prediction accuracy to “decode” the complexity of the brain (Grootswagers et al., 2017). These methods are powerful exploratory approaches that can explain a large amount of cross-validation variance and can be used for hypothesis testing when constrained in experiments. Yet a Bayesian model, in which every parameter is more deliberately chosen and interpretable, can act as a specific testable hypothesis more directly linked to a theory. In modern computer science research, the distinction between machine learning and Bayesian statistics is blurry as machine learning software such as TensorFlow can estimate posteriors (Štrumbelj et al., 2024), or alternatively Bayesian principles are often used for deep learning algorithms and it seems likely that the most effective tools will use a combination of the approaches (Papamarkou et al., 2024). Thus similarly, the field of neuroimaging may want to use the complimentary innovations of machine learning and Bayesian statistics, as opposed to relying on interpreting decoding prediction accuracy alone, as it has been suggested to lead to misinterpretations and irreproducible findings (Hullman et al., 2022).

The model results highlight some known qualities of the ssVEP measure, as well as some new details possibly related to the simultaneous fMRI recording. The first obvious trend in the data is a steady decrease in the overall ssVEP amplitude over trials which we term here adaptation (Figure 1). We found that there was strong evidence for an average decrease present in nearly all participants (Supplemental Figure 1; intercept and fatigue posteriors at https://osf.io/hfbjp/). It is known that the ssVEP adapts normally with repeated exposures (Norcia et al., 2016). However, the decrease in the present study is much more pronounced than has been seen in previous conditioning studies (McTeague et al., 2015; Wieser et al., 2014). The most obvious difference between this and previous studies is that participants are laying down with simultaneous fMRI acquisition. Thus, adaptation could be accentuated by heightened sympathetic arousal at the beginning of the study (because of the loud MRI scanner) and lower arousal at the end when compared previous EEG studies in which participants are seated in a chair. This could be represented in changes in the total global field power that may influence the recoverable ssVEP. Future research will model this spectral information, with a particular focus on alpha-band oscillation changes, to see how they are related to the ssVEP amplitude. However, the effect could also be partly explained by a physical interaction between the EEG system and MRI. The EEG-system is not immune to effects of the MRI and heats up during the study at the sensors and amplifier (Egan et al., 2021). Future research will examine if some aspect such as the gain or resistance recorded during the scan can be used to explain the adaptation effect. If these factors are present, they can be modeled explicitly in the present Bayesian framework. Lastly, it is known that a certain percentage of participants will not show an ssVEP amplitude when exposed to a flickering stimulus (Moratti et al., 2007; Regan, 1989). This is likely another reason why the models do not fit well for some participants. The signal to noise ratio of the ssVEP driving frequency compared to surrounding frequencies could be used to assess the presence of an ssVEP once the other confounding factors mentioned are addressed. If a participant does not have a ssVEP, then for that participant the ssVEP should not be correlated with alpha-oscillations, heart-rate changes, or fMRI. Some threshold, which could be determined in a data-driven portion of the model to decide at a per-participant basis if the ssVEP is informative for these other physiological measures.

The present project puts into practice the suggested Bayesian workflow (Gelman et al., 2020; Gabry et al., 2019; Schad et al., 2021). This is the process of iterative model building, comparison, visualization, and analysis that coincides with a formalizing and sharpening of scientific theories. Here, this involved comparing models on ELPD to gauge performance, and then breaking down the models to see how they differ in their accuracy and predictions. In the present project, the interpretation was straightforward as the learning model matched the competing models or did better on all metrics. Yet, a detailed look at the model makes apparent aspects that could be improved on in the future. Future work will compare competing models of learning (e.g., Pearce-Hall or Mackintosh models; Roesch et al., 2012) with neurophysiological informed mechanisms of attention and vision interactions (Desimone & Duncan, 1995; Reynolds & Heeger, 2009) to explain concurrently recorded EEG-fMRI data. Systematic model building and comparison will be necessary to assess the complexity of these interlocking assumptions.

The present learning model was kept relatively simple for interpretability, and there are several aspects that could be improved or expanded upon in future work. The first aspect was poorer fit in the habituation phase compared to Model 2. It is possible that the orientations of different cues may naturally produce a slightly larger amplitude at different scalp locations. Thus, fitting an intercept per cue may be justified. However, it can be seen in the supplemental figures that there were a variety of outliers in the habituation phase, which is a likely reason Model 2 looks better during that early phase of the study because it had the flexibility to accommodate them. Separately, the adaptation slope may not be an independent or linear relationship. It may be related to the gain of the equipment used and thus could be explained quantitatively. Even so, it may not be linear and could flatten / plateau over the course of the study. A multilevel Gaussian process regression or a spline function would allow for the modeling of adaptation to fit this phenomenon. Continuing, the scaling of CS+ associate strength per cue was done via an array per cue which did not allow for the pattern of generalization to differ between participants. Future work could combine this scaling across cues via a multilevel Ricker or Wavelet function that can fit generalization and sharpening patterns (Ahumada et al., 2025). This may improve statistical power by reducing the number of parameters to make predictions per cue. Lastly, the learning rates found suggest that the Rescorla-Wagner may not be the best option for predicting learning for the ssVEP. The ssVEP has been tied more explicitly to attention than other physiological responses (Müller et al., 1998, 2000, 2003). Although learning rates could be anywhere between 0 and 1, they concentrated at the two extremes. This is why the inverse logit transformed version of the model most likely fit better than the alternative learning models, as it is a transform that is primarily used to aid in fitting binary outcomes of a logistic regression. The original Rescorla-Wagner model would also force the use of one learning rate for paired and unpaired trials, as theoretically only the surprise related to prediction error should influence updates in learning. Thus, a different all-or-nothing model of learning may be better suited. Other experimental factors could have influenced this quick learning such as the long habituation phase and initial 100% pairing for the first 6 CS+ trials. This could have allowed participants to more easily differentiate the patterns and learn the relationship as other studies have found more of a gradual increase across trials (Santos-Mayo et al., 2025). Nonetheless, it is informative to know such a quick learning rate can occur for the ssVEP. Other physiological measures like skin conductance or pupil diameter may take longer to be conditioned as they perhaps involve a slower process of learning.

## Conclusion

This technical report sought to validate a multilevel learning model for analyzing ssVEP amplitudes during aversive conditioning, using innovations in Bayesian estimation algorithms and model comparison. While the learning model was relatively complex, the multilevel structure led to superior cross-validation accuracy compared to simpler models. Going beyond previous research that used single-trial estimates to draw meaning from a limited number of parameters, the present approach is a theory-driven generative model providing granular and reliable estimates at the level of individual subjects, cues, and trials. Thus, it was possible to visualize individual learning rates, the associate strength between the CS+ cue and noxious US, and how this predicted ssVEP amplitudes per cue and trial. Continuing, the conditioning effects per cue were better recovered than Model 2 that held the effect of cue constant between participants. Intrinsic to this workflow is the transparency and specificity of model results, which quantify the per-participant statistical certainty of meaningful factors such as the adaption of the ssVEP over trials and cross-validation variance explained. The findings provide a detailed understanding towards future model building efforts. Lastly, because the approach is a generative model, it was shown how missing trials can be interpolated based on the entire model’s structure. This may prove essential for multimodal physiological paradigms in which simultaneously recorded measures—each which a different collection of missing trials—are fused together. Thus, Bayesian workflows as illustrated here may serve as the theoretical basis for more ambitious future work, fusing not only different imaging methods, but integrating and contrasting more complex theories of visual, attention, and aversive learning.

## Supplemental materials

The code for preprocessing, models, analyses, figures can be found on the OSF page (https://osf.io/hfbjp/). The Farkas_Gaborgen_EEG_method_paper_equations.html details the rationale for prior selection and information on the 5 auxiliary models fit but not presented in the manuscript. Moving this information to a supplemental document was done to enhance transparency while keeping the manuscript readable. The statistical results for the adaptation decrease is presented below in Figure S1. There, 100 posterior draws are shown for the average adaptation as well as the median per participant per model. The median is shown per participant to aid in visualizing the difference between-participants. The full posterior adaptation intercept and slope per model can be found on the OSF page.

**Supplemental Figure 1.**
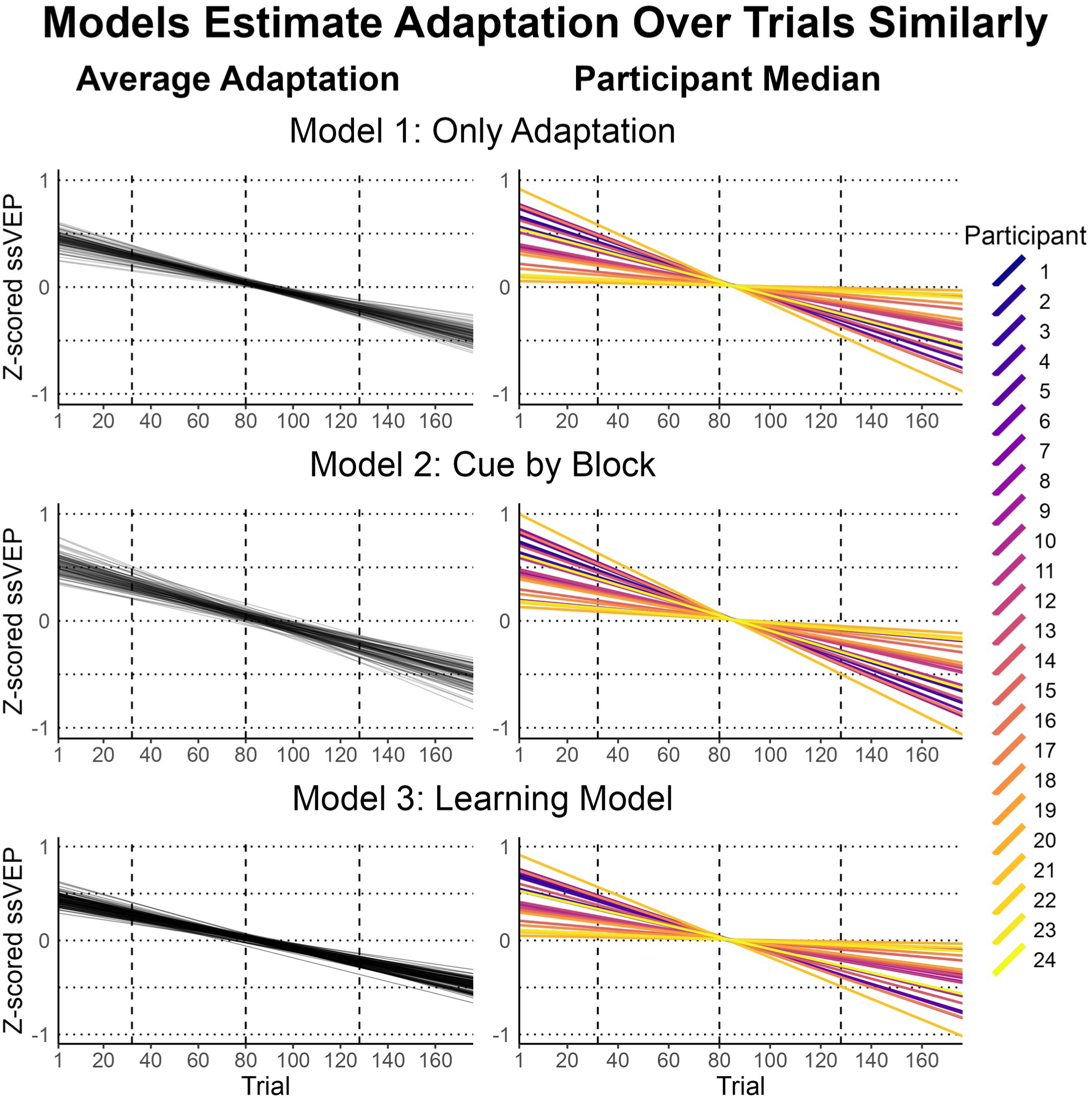
A visualization of the adaptation posterior per model. Left: 100 posterior draws of the average adaptation. The average adaptation posteriors are used as the multilevel prior for each participant informing and regularizing the right plots. Right: for interpretability, only the median posterior draw is of each participant. All three models found strong evidence for a downward trend in the ssVEP, which was similar between participants.

## Notes

https://osf.io/hfbjp/

### Competing Interest Statement

The authors have declared no competing interest.

